# Balancing cell populations endowed with a synthetic toggle switch via adaptive pulsatile feedback control

**DOI:** 10.1101/851212

**Authors:** A. Guarino, D. Fiore, D. Salzano, M. di Bernardo

## Abstract

Controlling cells endowed with the genetic toggle switch has been suggested as a benchmark problem in synthetic biology. It has been shown that a carefully selected periodic forcing can balance a population of such cells in an undifferentiated state. The effectiveness of these control strategies, however, can be mined by the presence of stochastic perturbations and uncertainties typically observed in biological systems and is therefore not robust. Here, we propose the use of feedback control strategies to enhance robustness and performance of the balancing action by selecting in real-time both the amplitude and the duty-cycle of the inducer molecular signals affecting the toggle switch behavior. We show, via in-silico experiments and realistic agent-based simulations, the effectiveness of the proposed strategies even in presence of uncertainties and stochastic effects. In so doing, we confirm previous observations made in the literature about coherence of the population when pulsatile forcing inputs are used but, contrary to what proposed in the past, we leverage feedback control techniques to endow the balancing strategy with unprecedented robustness and stability properties. We compare via in-silico experiments different control solutions and show their advantages and limitations from an in-vivo implementation viewpoint.

## Introduction

The Genetic Toggle Switch, implemented in-vivo for the first time by Gardner and Collins (*1*), has been highlighted as a fundamental synthetic circuit to endow cells with memory-like features (*2*) or to differentiate mono-strain cultures into different populations (*3*–*6*). A crucial problem in all reversible bistable systems is to reset their state by means of appropriate inputs to balance the system in an indeterminate state located in between the two stable states. A striking example is dedifferentiation in stem cells applications (*7*, *8*) where a terminally differentiated cell reverts and is maintained into an undifferentiated stem cell type.

As a paradigmatic example, we consider here the problem of balancing the genetic toggle switch (*1*) in a region surrounding its unstable equilibrium by manipulating two external inputs, the inducer molecules aTc and IPTG, that can affect its dynamics, see Fig. 1a). Solving this problem was highlighted as an important benchmark (*9*) in the applications of control theory to synthetic biology, similar to that represented by the classical inverted pendulum stabilization in control engineering (*10*).

**Figure 1:**
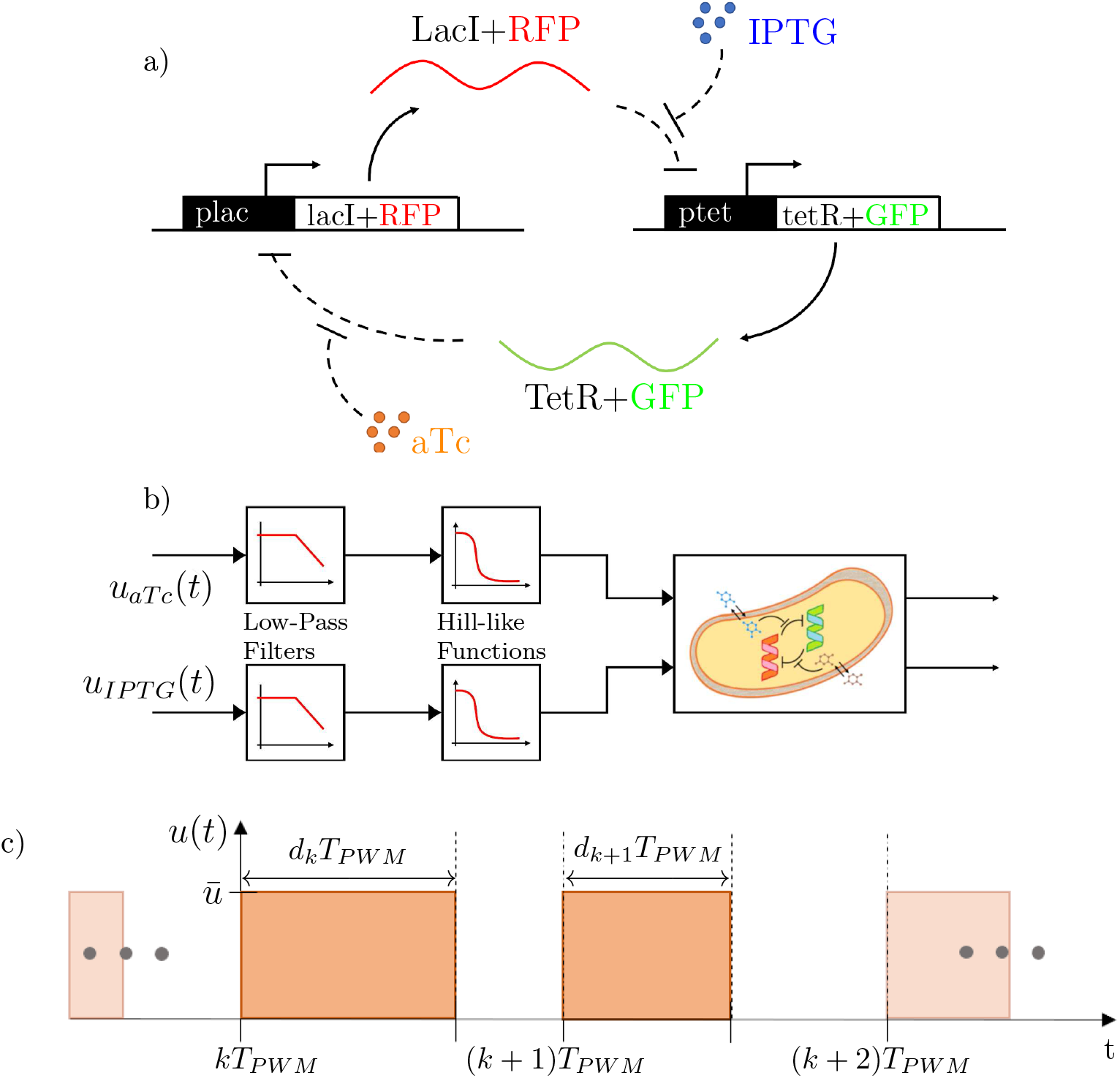
**Panel a)** <u>Schematic of the Genetic Toggle Switch network structure</u>. The two genes (LacI and TetR) ‒respectively bound with RFP(mKate2) and GFP(mEGFP)–mutually repress each other; the external inducers, IPTG and aTc, modulate the strengths of the repression exerted by LacI and TetR on each other. **Panel b)** <u>The toggle switch as a multi-input, multi-output (MIMO) control system</u>. The control inputs are first filtered because of diffusion through the cell membrane and then enter the toggle switch nonlinearly through Hill-like functions. **Panel c)** <u>Sketch of one of the pulsatile inputs applied to control the toggle switch</u>. Being mutually exclusive, the other input is chosen as its mirror image. *T*_*PWM*_ is the period of the inputs, while *d*_*k*_ represents the fraction of the *k*-th period during which the input is switched ON (the other being OFF).

Recently, it was shown that pulsatile periodic inputs with carefully selected periods, duty-cycle and amplitudes can be used to effectively balance a population of cells endowed with a genetic toggle switch (*9*) so that cells remain in a region where neither of the two genes is fully expressed. Moreover, when such forcing is used, high coherence was observed both in-silico and in-vivo across cells in the population.

The problem is that such desirable effects were only observed for certain forcing inputs whose characteristics (amplitudes, period, and duty-cycle – see Fig. 1c)) had to be carefully selected off-line by trial-and-error in order to achieve the desired balancing goal. Therefore, it was noted that when different periods and amplitudes were tested in-vivo, often coherence and control were lost with many cells in the population falling towards one of the two stable equilibria characterizing the switch rather than remaining balanced onto the desired undifferentiated state. Moreover, cell growth, cell-to-cell variability, uncertainties and noise can render any off-line choice of the period and amplitude unable to reach the control goal in practice. Therefore, there is the need of synthesizing a closed-loop action able to compensate against these effects by adapting inputs’ features in real-time to cope dynamically with changing environmental conditions, growth, diffusion and other unmodelled effects.

Previous studies about the time-scale of the filtering properties of the cell membrane (*11*, *12*) suggest that an appropriate choice of period of the forcing inputs is about 240 minutes. The goal of this paper is, then, to propose a new strategy based on feedback control to select and adapt in real-time the features of the input signals (amplitudes and duty-cycle) while maintaining their periodic nature. This allows us to exploit the beneficial effects of periodic forcing inputs (*9*) while endowing the population with stability, coherence and robustness (*13*) that cannot be achieved with open-loop approaches such as those previously presented in the Literature. The challenge from a theoretical view point is to achieve this goal by means of two mutually exclusive inducer molecular inputs that can be provided to all the cells, whose duty-cycle can be varied in real-time as a function of the measured cell population behavior.

Other control approaches to steer the behavior of a genetic toggle switch include Pulse-Shaping Control (*14*–*16*) and Reinforcement Learning (*17*) to drive the switch from one stable state to the other, or Stochastic Motion Planning (*18*) and Piecewise Linear Switched Control (*19*, *20*) that were used in-silico to stabilize a toggle switch about its unstable equilibrium. Differently from previous results in the Literature, our theoretical approach is strongly oriented to the assessment of a possible in-vivo implementation of the control strategies being presented, taking into account for their validation realistic constraints on the inputs and other phenomena such as cell growth and diffusion, as described later in this work. Our results complement and integrate those presented by Lugagne et al. (*9*) that were the first to report in-vivo control experiments involving a population of cells endowed with the genetic toggle switch.

## Results and Discussion

We focus on designing control strategies to stabilize the toggle switch implementation shown in Fig. 1a), using the model equations parametrized from experimental data that were derived recently in (*9*) and further analyzed in our previous work (*11*) (see Sec. 1 of the SI for more information and details). From a control viewpoint, the effect of the control inputs can be summarized as in Fig. 1b) where we see that the inducer molecules are filtered by the cell membrane and then act onto the switch dynamics through nonlinear (Hill-like) time-varying input functions in the model.

Firstly, we tested the effect of applying the inputs in an open-loop fashion by using pre-computed amplitudes, say *ū*_*aTc*_, *ū*_*IPTG*_, and duty-cycle, say *d*_*k*_, as in the implementation proposed by Lugagne et al. (*9*). We found that, as shown in Fig. 2, the strategy fails to achieve the balancing goal in the presence of diffusion, cell-to-cell variability and noise. Therefore, to balance the toggle switch population, we then synthesized a feedback (closed-loop) control approach that is still based on using two mutually exclusive periodic inputs but is able to adapt and change their duty-cycle and select their amplitudes to better achieve the desired control goal.

**Figure 2:**
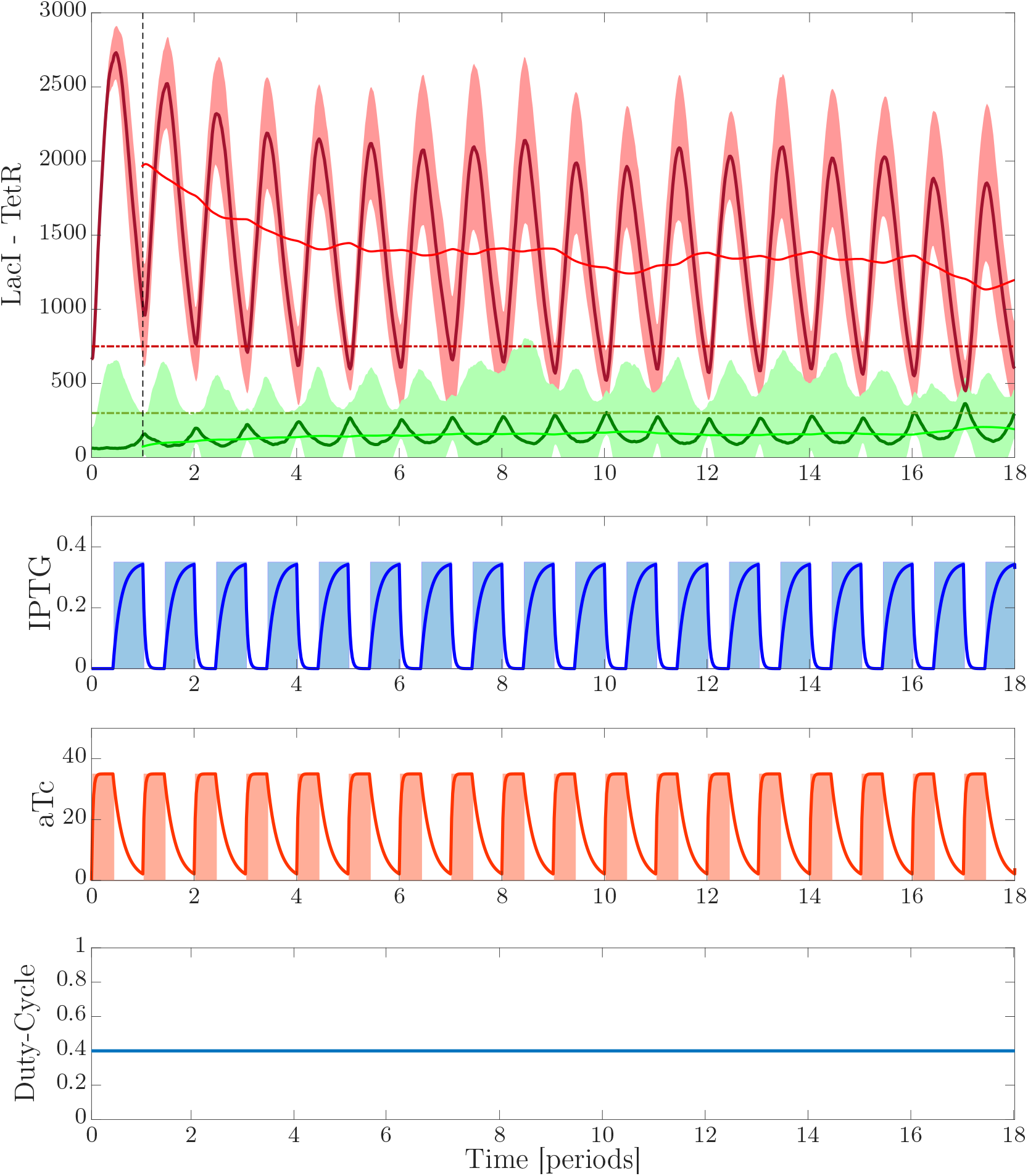
<u>Open-loop periodic forcing</u>. The evolution of TetR and LacI is shown when cells are subject to mutually exclusive pulsing inputs whose amplitude and duty-cycle were pre-computed off-line to be *ū*_*aTc*_ = 35, *ū*_*IPTG*_ = 0.35, *d*_*k*_ = *d*_*ref*_ = 0.4. The period was fixed to *T*_*PWM*_ = 240 min. Total simulation time shown is 72 hours. Population size: 17 cells. Top panel: Dashed red and green lines are the desired setpoints, set respectively to *LacI*_*ref*_ = 750 and *TetR*_*ref*_ = 300. Solid dark red and green lines are the evolution of the average values over the population of *LacI* and *TetR* as expressed by the cells during the experiment. Shaded areas represent the value of the standard deviation from each mean value, computed at each time instant. Solid light red and green lines are the evolution of the mean value of the oscillations evaluated with a moving window of period equal to *T*_*PWM*_. The lack of convergence to the desired set-point clearly shows the limits of open-loop, pre-computed inputs in achieving the control objective when diffusion and other effects are appropriately modeled. Middle panels show the evolution of the pulsing inputs applied to the system. Bottom panel shows the the duty-cycle that is kept constant over time.

In particular, we adopted an external control strategy (*21*) that can be implemented in microfluidics via a fluorescence microscope and actuated syringes as depicted in Fig. 3a). The core of the strategy is the feedback control algorithm that is summarized in Fig. 3b). The strategy is based on two control actions: 1. A *feedforward* action that pre-computes the value of the input amplitudes and duty-cycle ideally required to achieve the control goal in the absence of perturbations and diffusion effects (this is done by inverting the simplified, average nonlinear model of the toggle switch subject to periodic inputs we derived in earlier work (*11*) – for further details see Methods); 2. A *feedback* action that adapts the duty-cycle *d*_*k*_ of the periodic inputs as a dynamic function of the current cell behavior, measured via the fluorescence microscope.

**Figure 3:**
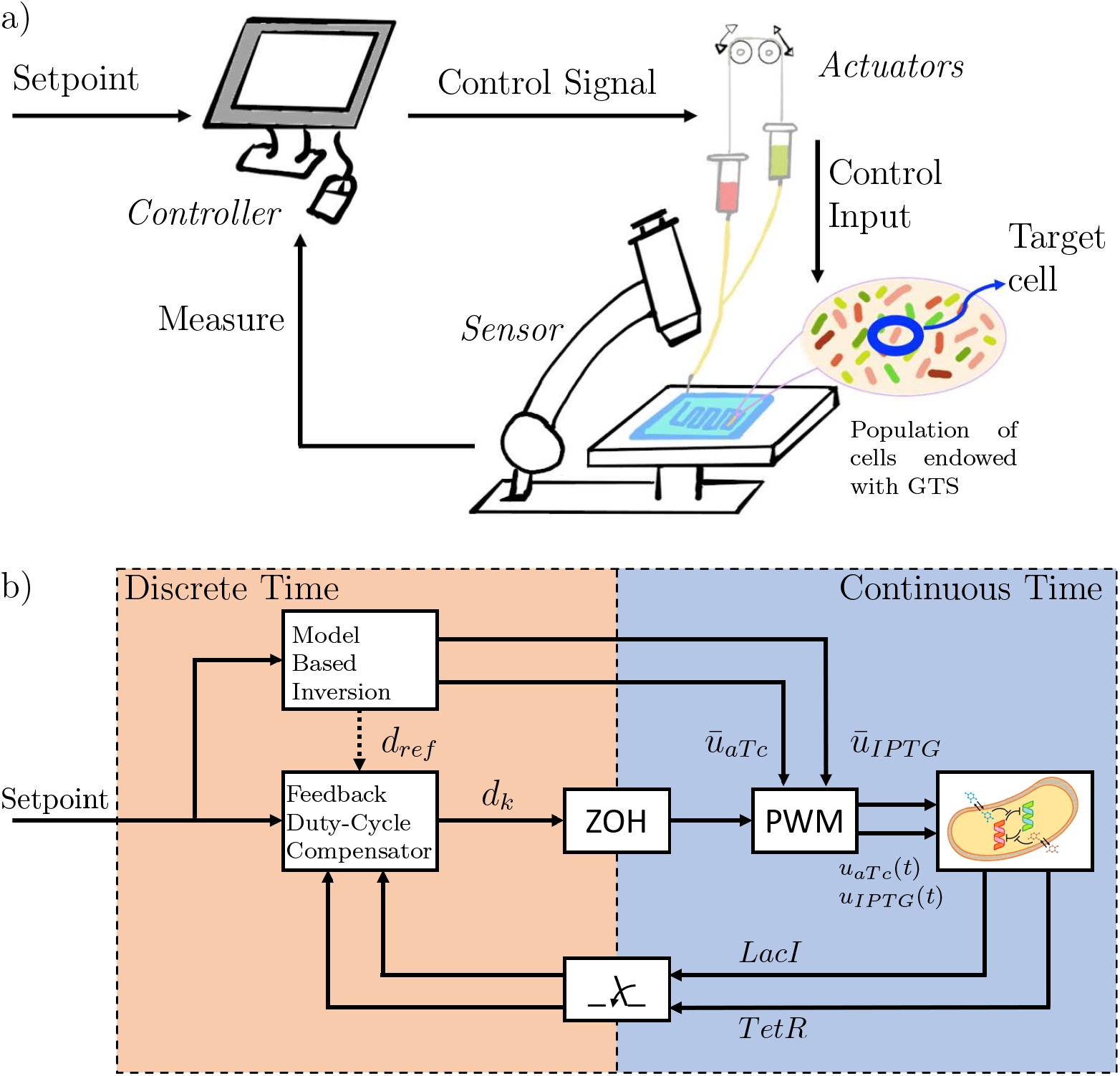
**Panel a)** <u>External Control Architecture</u>. A population of E. coli endowed with the Genetic Toggle Switch is hosted in a microfluidic device. A fluorescence microscope takes pictures of the cells, whose average RFP and GFP values are evaluated through segmentation algorithms. This information, together with the setpoint of the experiment, is sent to the controller that computes *online* the inputs to be applied to the cells. The actuators, a system of motorized syringes, receive the control signal and produce the action needed to feed the population of cells in the chamber with the required inputs. **Panel b)** <u>Block diagram of the proposed closed-loop hybrid control strategy</u>. The population of cells, together with the PWM inputs, evolve in continuous-time. The controller is instead designed in discrete-time, computing the control input at each time period *T*_*PWM*_. A feedforward *Model Based Inversion* block evaluates the amplitudes *ū*_*aTc*_ and *ū*_*IPTG*_ of the pulse wave inputs, on the basis of the setpoint [*LacI*_*ref*_ *TetR*_*ref*_]. The feedback controller evaluates and adapts in real time the duty-cycle of the inputs as a function of the desired setpoints and the (sampled) outputs of the system. The Zero-Order Holder (ZOH) keeps the duty-cycle, computed by the compensator at the beginning of each period, constant during the rest of the period *T*_*PWM*_.

We compare the effectiveness of two feedback control strategies: a proportional-integral (PI) controller that drives a pulse width modulation block (PI-PWM, see Sec. 4 of the SI for further information and details) and a Model Predictive Controller (MPC) that solves the problem by optimizing a desired cost function to select the input duty-cycle dynamically (see Sec. 5 of the SI).

Figure 4 and Figure 5 show the results of the deterministic and stochastic in-silico experiments we conducted. We see that both strategies are effective in controlling the toggle switch to the desired output value despite the presence of perturbations and uncertainties that render other approaches unreliable such as the open-loop strategy (*9*), shown in Figure 2, where the inputs are pre-computed off-line.

**Figure 4:**
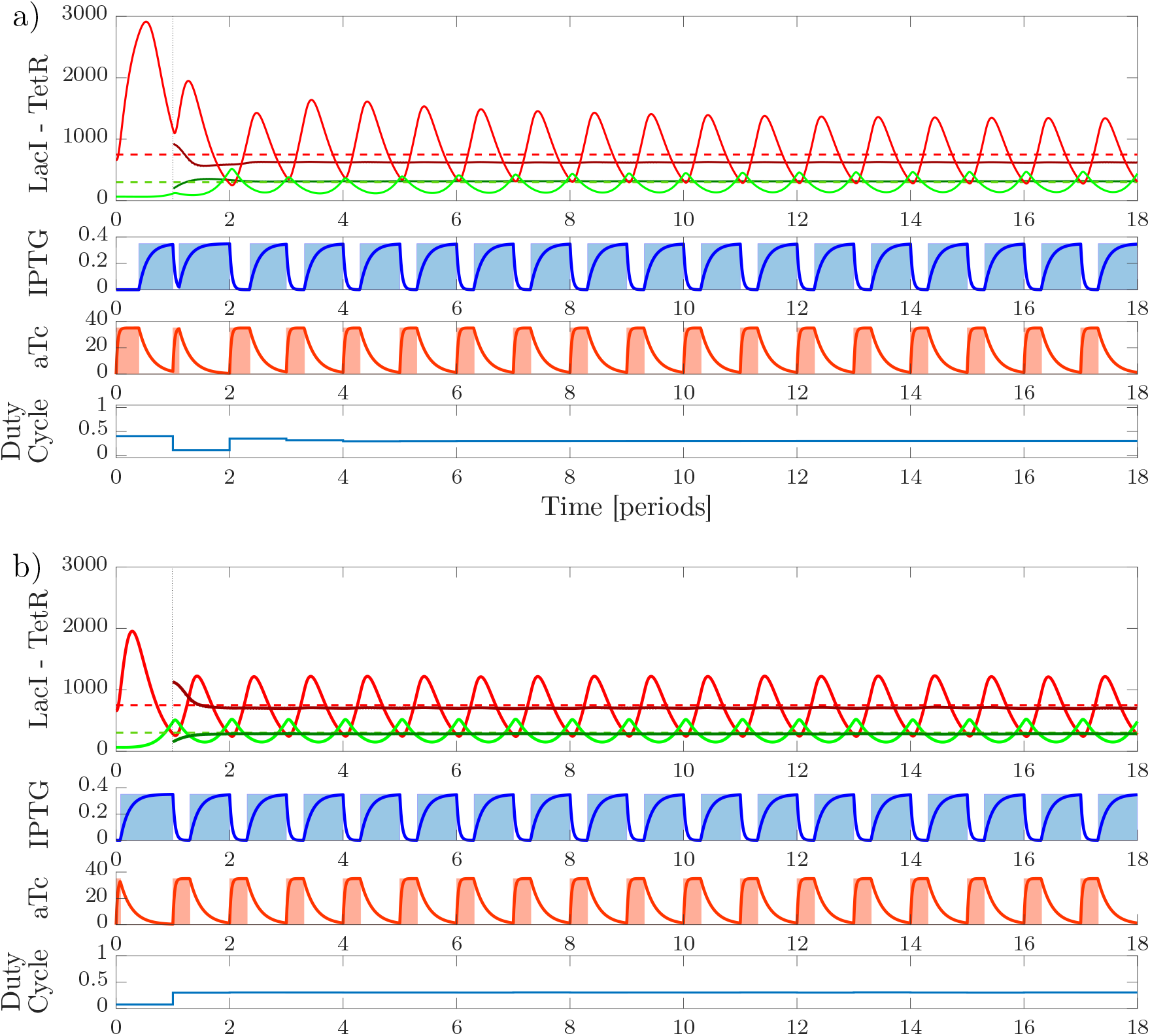
<u>In-silico deterministic experiment</u>: comparison of the performance of the PI-PWM (**panel a)**) and MPC (**panel b)**) control strategies via deterministic simulations. The value of the input amplitudes are set to *ū*_*aTc*_ = 35, *ū*_*IPTG*_ = 0.35. For the PI-PWM, the duty-cycle starts from *d*_*ref*_ = 0.4 and is then adapted by the controller after the first period, while the MPC computes the duty-cycle from the start (solving the optimization problem). PI gains were set to *k*_*P*_ = 0.0101 and *k*_*I*_ = 0.0401. The parameters of the MPC cost function parameters were set to *K*_LacI_ = 1, *K*_TetR_ = 4, while the prediction horizon is equal to 3*T*_*PWM*_. Total simulation time is 72 hours. Top panels: dashed red and green lines represent the setpoint of the experiment, respectively *LacI*_*ref*_ and *TetR*_*ref*_. Solid lines show the evolution of promoter proteins for *LacI* (red) and *TetR* (green). Dark solid lines, starting from *t* = *T*_*PWM*_, are the mean values of the state in the time period, evaluated with a moving window of period *T*_*PWM*_. Middle panels show the evolution of the pulsing inputs applied to the system. Bottom panels show the evolution of the duty-cycle over time.

**Figure 5:**
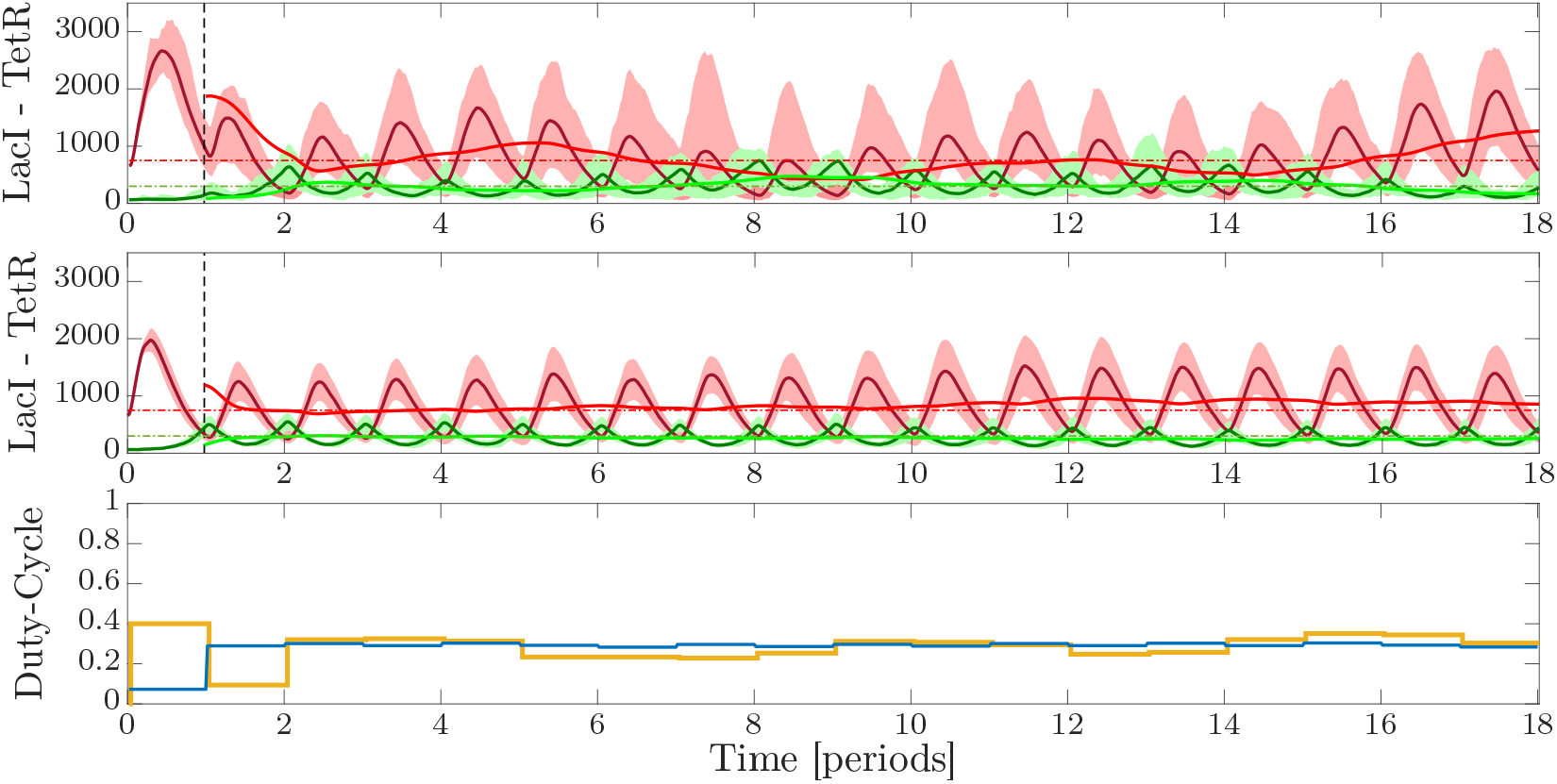
<u>In-silico stochastic experiment</u>: comparison of the performance of the PI-PWM (top panel) and MPC (central panel) control strategies. Inputs amplitudes are set to: *ū*_*aTc*_ = 35, *ū*_*IPTG*_ = 0.35 while *d*_*ref*_ = 0.4 (for the PI-PWM), *T*_*PWM*_ = 240 min. PI gains were set to *k*_*P*_ = 0.0101 and *k*_*I*_ = 0.0401. MPC’s cost function parameters have been set to *K*_LacI_ = 1, *K*_TetR_ = 4, while the prediction horizon is 3*T*_*PWM*_. Total simulation time is 72 hours. Population size: 17 cells. Dashed lines are the setpoint of the experiment, for *LacI*_*ref*_ (red) and *TetR*_*ref*_ (green). Solid red and green lines are the average evolution of *LacI* and *TetR* over the population. Darker solid lines represent the evolution of the mean trajectory in the period, evaluated with a moving window as in the deterministic case. Shaded areas represent the values of the standard deviation from the means, at each time instant. Bottom panel: Duty-cycle evolution over time when the MPC (blue) or the PI-PWM (yellow) strategies are used.

In general, we find that in the deterministic case the MPC – often used in control applications in synthetic biology (*22*) – guarantees better performance in terms of dynamic regulation; the overshoot and transient duration being significantly lower than those observed with the other strategy. Moreover, the MPC achieves also better steady-state regulation of the setpoint with respect to the PI-PWM controller over the 18 periods considered as realistic for the achievement in-vivo of the control goal. This is essentially due to the approximations made by the analytical average model and the projector on the (pre-computed) curves of equilibria used to compute the control inputs of the PI-PWM, which cause a residual error at steady-state (see Sec. 4 of the SI for further details). A quantitative assessment of the controllers’ performance is reported in Tab. S2 in Sec. 7 of the SI and confirms the qualitative observations made above. The stochastic simulations in Figure 5 show that, qualitatively, the results of the in-silico experiments conducted via deterministic numerical simulations still hold. However, the gap in performance between the strategies assessed quantitatively in Table 3 in the SI is reduced when stochastic effects are included in the simulation, with the MPC still showing better performance than the PI-PWM.

As a further validation of the effectiveness of these control strategies, we carried out robustness tests by introducing parametric variations in the cell population to model cell-to-cell variability. Specifically, the parameters of each cell in the population were independently drawn from Gaussian distributions centered on their nominal values given in Table 1 in the SI with a standard deviation of 5% and 10%, respectively. Quantitative metrics are reported in Table 4 in the SI. The results confirm that PI-PWM is more robust to small perturbations as it does not rely on exact knowledge of the model. In contrast, larger parameter perturbations worsen considerably the PI-PWM performance, making the adoption of MPC preferable for in-vivo implementation.

**Table 1:** Value of the parameters of the Model (1)-(6) taken from (*9*).

|  |  |  |  |
| --- | --- | --- | --- |
| $\kappa_L^{m0}$ | 3.20e-2 mRNA min<sup>-1</sup> | $g_L^m, g_T^m$ | 1.386e-1 min<sup>-1</sup> |
| $\kappa_T^{m0}$ | 1.19e-1 mRNA min<sup>-1</sup> | $g_L^p, g_T^p$ | 1.65e-2 min<sup>-1</sup> |
| $\kappa_L^m$ | 8.30 mRNA min<sup>-1</sup> | $\theta_{LacI}$ | 31.94 a.u. |
| $\kappa_T^m$ | 2.06 mRNA min<sup>-1</sup> | $\eta_{LacI}$ | 2.00 |
| $\kappa_L^p$ | 9.726e-1 a.u. mRNA min<sup>-1</sup> | $\theta_{TetR}$ | 30.00 a.u. |
| $\kappa_T^p$ | 9.726e-1 a.u. mRNA min<sup>-1</sup> | $\eta_{TetR}$ | 2.00 |
| $k_{IPTG}^{in}$ | 2.75e-2 min<sup>-1</sup> | $\theta_{IPTG}$ | 9.06e-2 mM |
| $k_{IPTG}^{out}$ | 1.11e-1 min<sup>-1</sup> | $\eta_{IPTG}$ | 2.00 |
| $k_{aTc}^{in}$ | 1.62e-1 min<sup>-1</sup> | $\theta_{aTc}$ | 11.65 ng/ml |
| $k_{aTc}^{out}$ | 2.00e-2 min<sup>-1</sup> | $\eta_{aTc}$ | 2.00 |

The effects of the same perturbations in open-loop is also reported and discussed in Sec. 7 of the SI for the sake of comparison.

With a view towards the in-vivo implementation, we assessed via agent-based simulations the performance of both control strategies. To this aim we conducted in-silico experiments with BSim 2.0, an advanced agent-based simulator of bacterial populations (*23*, *24*) that is able to replicate realistic phenomena such as cell growth, spatial diffusion, cell-to-cell variability, and flush-out of the cells from the microfluidic chamber (see Methods for further details).

Figure 6 shows the results of the agent-based simulations confirming the effectiveness and viability of the strategies for in-vivo experiments. Supplementary videos MOV1 and MOV2 are available online and show the simulations of the experiments adopting the two different control strategies, PI-PWM and MPC respectively.

**Figure 6:**
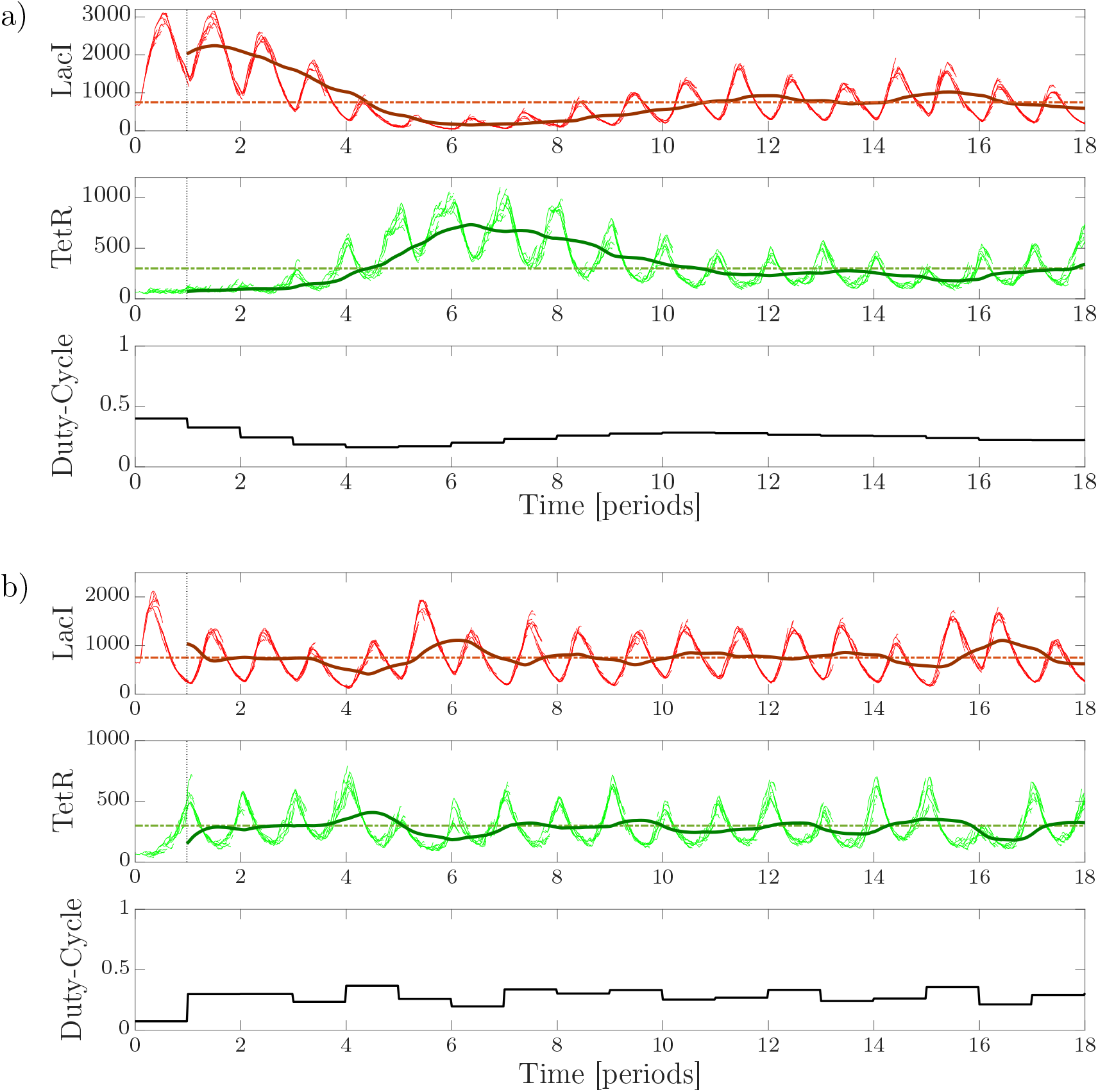
<u>Agent-based simulation in BSim 2.0</u> of the PI-PWM (**panel a)**) and MPC (**panel b)**) control strategies. The pulsatile inputs’ amplitudes were set to *ū*_*aTc*_ = 35, *ū*_*IPTG*_ = 0.35, while *d*_*ref*_ = 0.4 and *T*_*PWM*_ = 240 min. Total simulation time is 72 hours. We considered E. coli cells growing in a single chamber of a “mother machine”-like microfluidic device (*27*): the simulations start with a single cell located at the bottom of the chamber; as the cell grows and duplicates, it pushes outside of the chamber new cells that exceed the maximum capacity of the chamber (around 10 cells). The top panel shows the evolution over time of *LacI*; the dashed line representing the setpoint *LacI*_*ref*_ = 750, while lighter lines the evolution of the state for each cell in the simulation, and the darker solid line the mean trajectory computed over the population, evaluated through a moving window of period *T*_*PWM*_. The middle panel shows the evolution over time of *TetR*; the dashed line representing the setpoint *TetR*_*ref*_ = 300, lighter lines are the evolution of the state for each cell in the simulation, while the dark solid line represents the evolution of the mean trajectory across the population in the period, evaluated using a moving window of period *T*_*PWM*_. The bottom panel shows the evolution of the duty-cycle over time.

Finally, as previously done with stochastic simulations, we introduced a 10% variation of all parameters of each cell in the agent-abased model to reproduce cell-to-cell variability. The quantitative analysis reported in Tab. 5 of the SI confirms that the adoption of the MPC control strategy is to be preferred in the presence of such uncertainties.

To conclude, Model Predictive Control has been shown to guarantee consistently better control performance in all in-silico experiments carried out. However, its main limitation is the higher computational cost required to solve the optimization problem at the beginning of each control cycle and therefore requires an experimental platform with adequate computing power for its in-vivo implementation.

## Discussion

Starting from the observations that pulsatile periodic inputs can balance a toggle switch population in a region where neither of the two genes is fully expressed (*9*), we presented a suite of feedback control strategies that can be used to change and adapt the duty-cycle of the inputs in real-time and select their amplitudes to achieve robust stabilization of the population even in the presence of noise and other unavoidable effects which render previous open-loop approaches unviable. We demonstrated the ability of the proposed strategies to solve the balancing problem by a combination of deterministic and stochastic in-silico experiments and agent-based simulations under realistic assumptions on physical and technological constraints of a possible microfluidics experimental platform.

Our results show that using pulsatile inputs computed online by means of a feedback control strategy is a viable approach to achieve the robust stabilization of a toggle switch population about its undifferentiated state. Our findings are further supported by recent observations that pulsatile inputs are often exploited in place of constant ones in the regulation of biological processes (*25*, *26*).

We wish to emphasize that the control strategies we propose can be of practical relevance to “reset” other bistable, or multi-stable, cellular systems by balancing them in an unstable region corresponding to some undifferentiated state of interest. This is important, for example, in multicellular control experiments (*5*) where it has been proposed that monostrain populations can be differentiated into multiple subpopulations by flipping the state of a synthetic toggle switch associated to different functions, or in stem cell applications where dedifferentiation (*7*, *8*) aims precisely at “resetting” a differentiated cell.

From a theoretical viewpoint, we wish also to highlight that nonlinear average models capturing the system behavior under external pulsing stimuli could be useful for the design of feedback control strategies for future applications in synthetic biology.

## Methods

### In-silico Control Experiments

**Deterministic Simulations** have been conducted in MATLAB using the model derived by Lugagne et al. described in Sec. 1 of the SI. The numerical integration of the model has been carried out using an event-driven algorithm we developed. Each event is associated with the change of the inputs given to the system; therefore, an event is determined by the duty-cycle *d*_*k*_ and the period *T*_*PWM*_ of the PWM inputs which is set to 240 min. The solver ode45 generates 100 non uniformly distributed time samples in each time period of the inputs *T*_*PWM*_, leading to 1800 time samples in a total simulation time of 18 periods, corresponding to 4,320 min.

**Stochastic Simulations** have been conducted in MATLAB using the well-known Gillespie’s Stochastic Simulation Algorithm^1^. The solver is set at a fixed time step of 5 minutes. Using this setup, we obtain 48 time samples in each period *T*_*PWM*_, leading to 864 samples in a total simulation time of 18 periods. Multiple cells stochastic simulations have been conducted in parallel, using the MATLAB Parallel toolbox to speed up the computation.

**Agent-Based Simulations** have been conducted in BSim 2.0, an advanced bacteria simulator developed in Java (*23*, *24*). Inspired by the so-called *mother machine* (*27*), we designed a 1 × 30 × 1 *μ*m rectangular chamber that hosts a single layer cell population where cells are lined up. The chamber is open on the top (short side), from where the inducers diffuses and cells are flushed out due to their own growth and medium flow. The solver is based on the *Euler-Maruyama* (*28*) method and generates samples at a fixed time step of 5 minutes, leading to 48 samples per period and 864 samples in the total simulation time of 18 periods (Supplementary Videos 1 & 2 available online).

### Feedforward Model Based Inversion

To select the amplitude of the PWM inputs (and the nominal value of the duty-cycle for the PI-PMW strategy), we calculated a database of 60 curves of equilibria for our system (see Sec. 3 of the SI), using the time-averaged model of the circuit under periodic inputs we previously derived in (*11*). The average model is reported and described in Sec. 2 of the SI.

Using the model, we computed each curve of equilibria in the database considering 18 points per curve each representing the steady-state behavior of the time-averaged model of the toggle switch when the amplitudes of the two inputs *ū*_aTc_ and *ū*_IPTG_, and the nominal value of the duty-cycle *d*_*ref*_ are varied. Given the desired setpoint 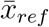 we query the database to find the equilibrium point 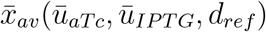 closest to it in terms of Euclidean distance. The values of the amplitudes – *ū*_*aTc*_ and *ū*_*IPTG*_ – and the duty-cycle *d*_*ref*_ that correspond to the closest equilibrium point 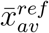 are then selected and used to implement the feedforward action in the control experiment.

### Feedback control strategies implementation

#### PI-PWM

The PI-PWM control technique evaluates the correction *δd*_*k*_, at every time instant *t*_*k*_ = *k T*_*PWM*_, that has to be added to the nominal duty-cycle *d*_*ref*_. Thus, the duty-cycle is evaluated online as follows:

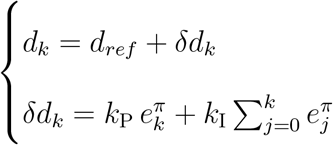

where *k*_P_ and *k*_I_ are the gains of the PI controller and 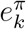 is the error measure computed using a nonlinear projector. For further information see Sec. 4 of the SI.

#### MPC

Model Predictive Control explicitly evaluates the duty-cycle *d*_*k*_ that minimizes the following cost function:

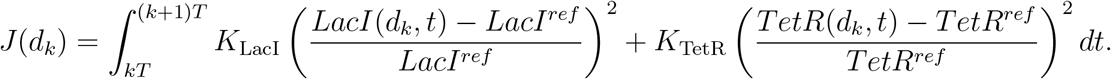

The optimization problem takes into account the constraints on the duty-cycle – whose value is bounded between 0 and 1 – and on the dynamics of the system. At the beginning of each period *t*_*k*_ = *k T*_*PWM*_, the states *LacI* and *TetR* of the system evolve according to the deterministic model (see Sec. 1) until the end of the prediction horizon *t*_*f*_ = (*k* + *n*_*ph*_)*T*_*PWM*_, where 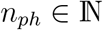 represents the number of periods *T*_*PWM*_ in the prediction horizon.

The optimization problem described above was solved using the *genetic algorithms* toolbox available in MATLAB^2^. The genetic algorithm is initialized with a random population of 50 individuals. The maximum number of stall generations was set to 30.

More details on the control strategy are available in Sec. 5 of the SI.

## Supporting information

MOV1

MOV2

## Associated Content

Supporting Information available. Equilibrium curves database available.

## Author Information

The authors declare no competing interests.

## Acknowledgments

The authors wish to acknowledge support from the research project COSY-BIO (Control Engineering of Biological Systems for Reliable Synthetic Biology Applications) funded by the European Union’s Horizon 2020 research and innovation programme under grant agreement No 766840. DS is supported by a University of Naples Federico II doctoral fellowship. AG wishes to thank Giansimone Perrino (Telethon Institute for Genetics and Medicine, Naples, Italy) for the helpful discussions on the implementation of MPC for biological systems.

## Author Contributions

AG implemented the controllers, carried out the simulations and analyzed the data. DF derived the mathematical model used for control design and contributed to the synthesis of the feedback controllers and the data analysis. DS and AG contributed to the controllers’ validation and implemented the agent-based study in BSim. MdB conceived the idea of the study. AG, DF and MdB wrote the paper.

## Supplementary Information

### 1 Model

The model of the synthetic toggle switch we considered in our analysis was originally developed in Lugagne et al. (*9*). The model captures the pseudo-reactions describing transcription

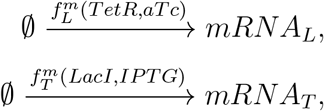

those describing translation

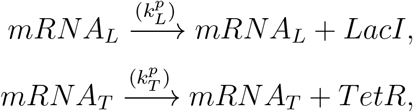

and those related to dilution/degradation

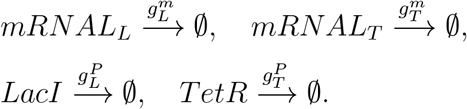

The pseudo-reactions listed above can be put together to obtain the following model:

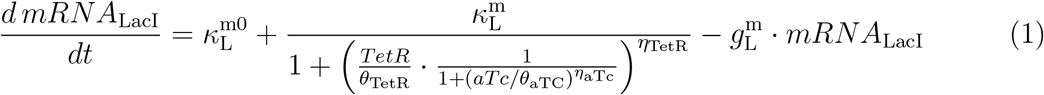

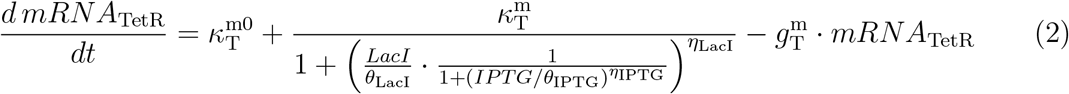

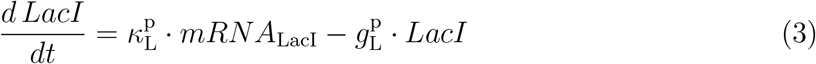

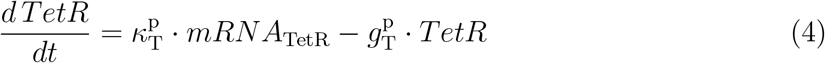

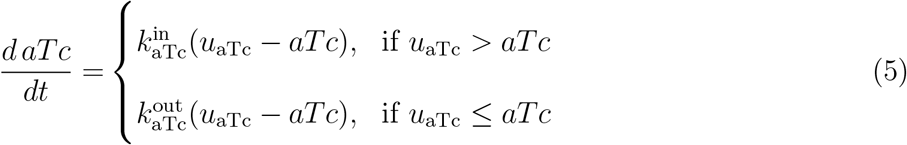

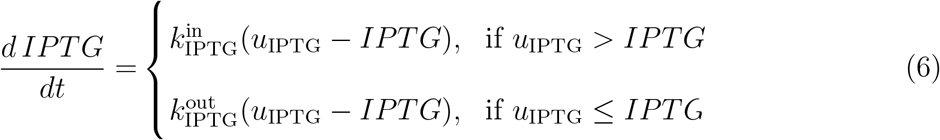

The parameters of the model are listed in Table 1.

### 2 Average Model

Since the time scales of the mRNA dynamics is notably faster than that of the proteins (*11*), by using quasi-steady state arguments, we can obtain the non-dimensional Quasi-Steady State Model:

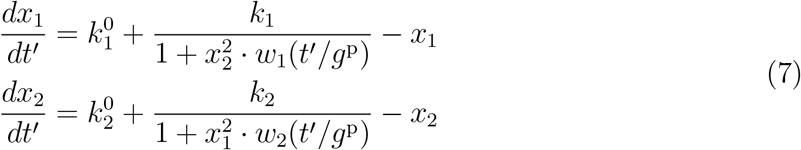

where

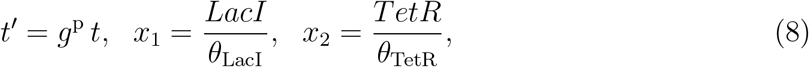

and the dimensionless parameters are defined as

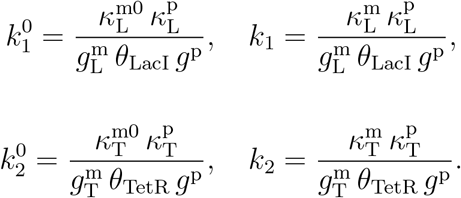

The inputs to the system are modeled by the nonlinear functions *w*_1_ and *w*_2_ whose expression was derived in our previous work (*11*).

When subject to mutually exclusive pulsatile inputs, of the form

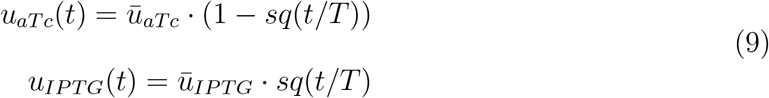

where *sq*(*t/T*) is a periodic square wave of period *T* and duty-cycle *d* that assumes values in [0, 1], model (7) yields the following *average vector field*:

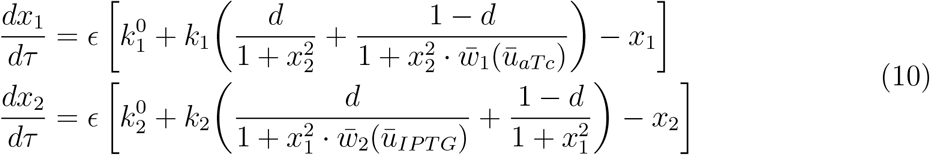

where *τ* = *t*′/*g*_*p*_*T*.

### 3 Curves of equilibria of the average model

The number and position in state space of the equilibrium points 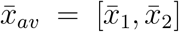 of the average vector field (10) depend on the specific choice of the amplitudes *ū*_*aTc*_ and *ū*_*IPTG*_ of the mutually exclusive pulsatile inputs, and on the value of the duty-cycle *d*. For example, for the reference values *ū*_*aTc*_ = 50 ng/ml and *ū*_*IPTG*_ = 0.5 mM, system (10) is monostable and the position of the equilibrium point 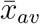 varies monotonically with *d* as reported in Figure 7 (blue dots). Hence, given certain values of *ū*_*aTc*_ and *ū*_*IPTG*_, it is possible to move the position of 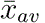 on the corresponding curve by varying *d*, as reported in Figure 8.

**Figure 7:**
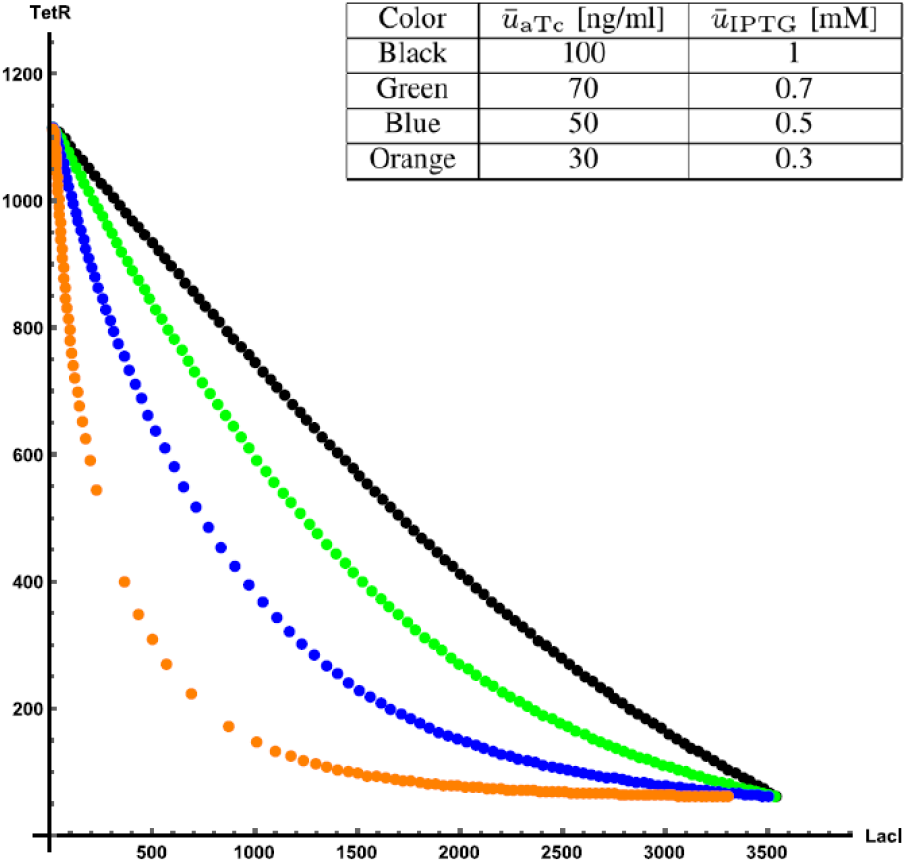
Equilibrium points 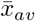 of (10) as a function of duty-cycle *d* re-scaled in arbitrary fluorescence units using (8). Each dot represents the location of the unique stable equilibrium point of system (10) evaluated for *d* taking values in the interval [0, 1] with increments of 0.01.

**Figure 8:**
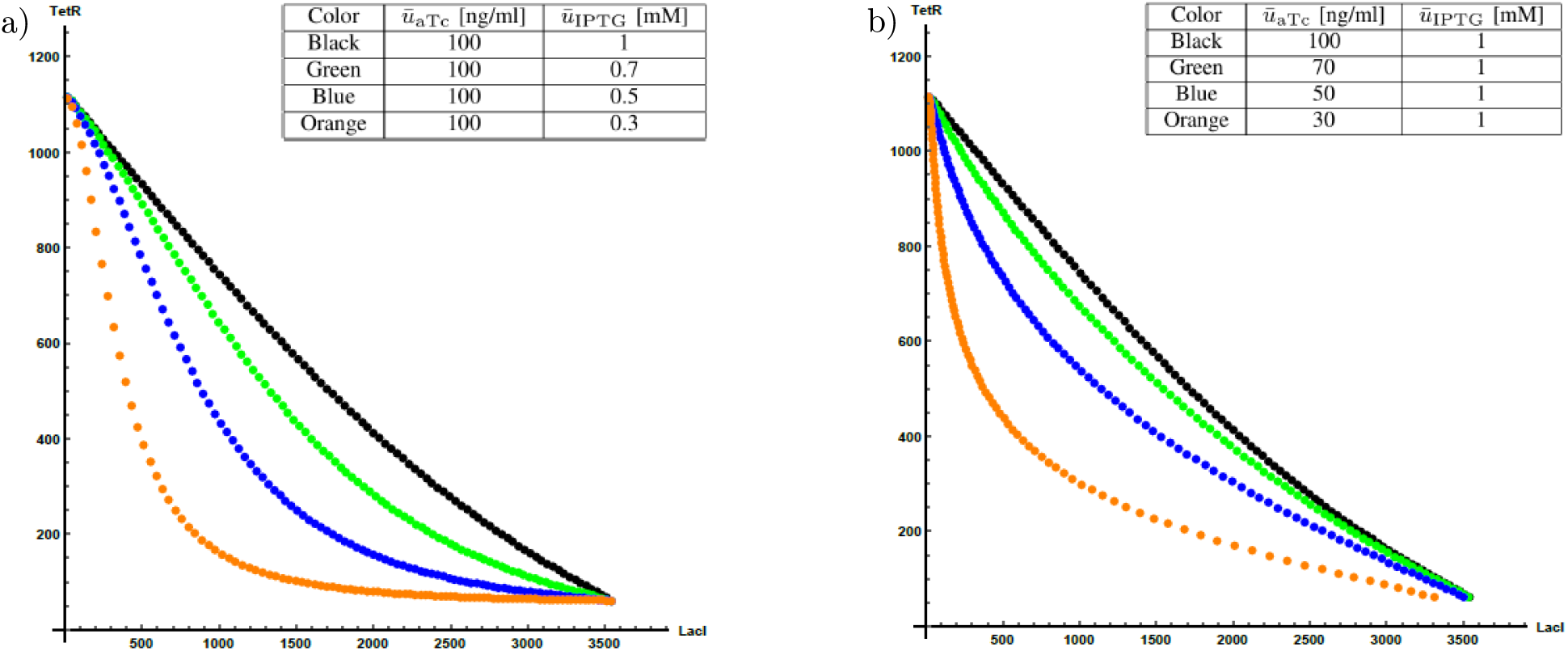
Equilibrium points 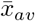 of (10) as a function of duty-cycle *d* re-scaled in arbitrary fluorescence units using (8). Each dot represents the location of the unique stable equilibrium point of system (10) evaluated for *d* taking values in the interval [0; 1] with increments of 0.01. **Panel a)**: Equilibrium points for *ū*_*aTc*_ = 100 ng/ml and different values of *ū*_*IPTG*_. **Panel b)**: Equilibrium points for 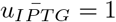 mM and different values of *ū*_*aTc*_.

**Figure 9:**
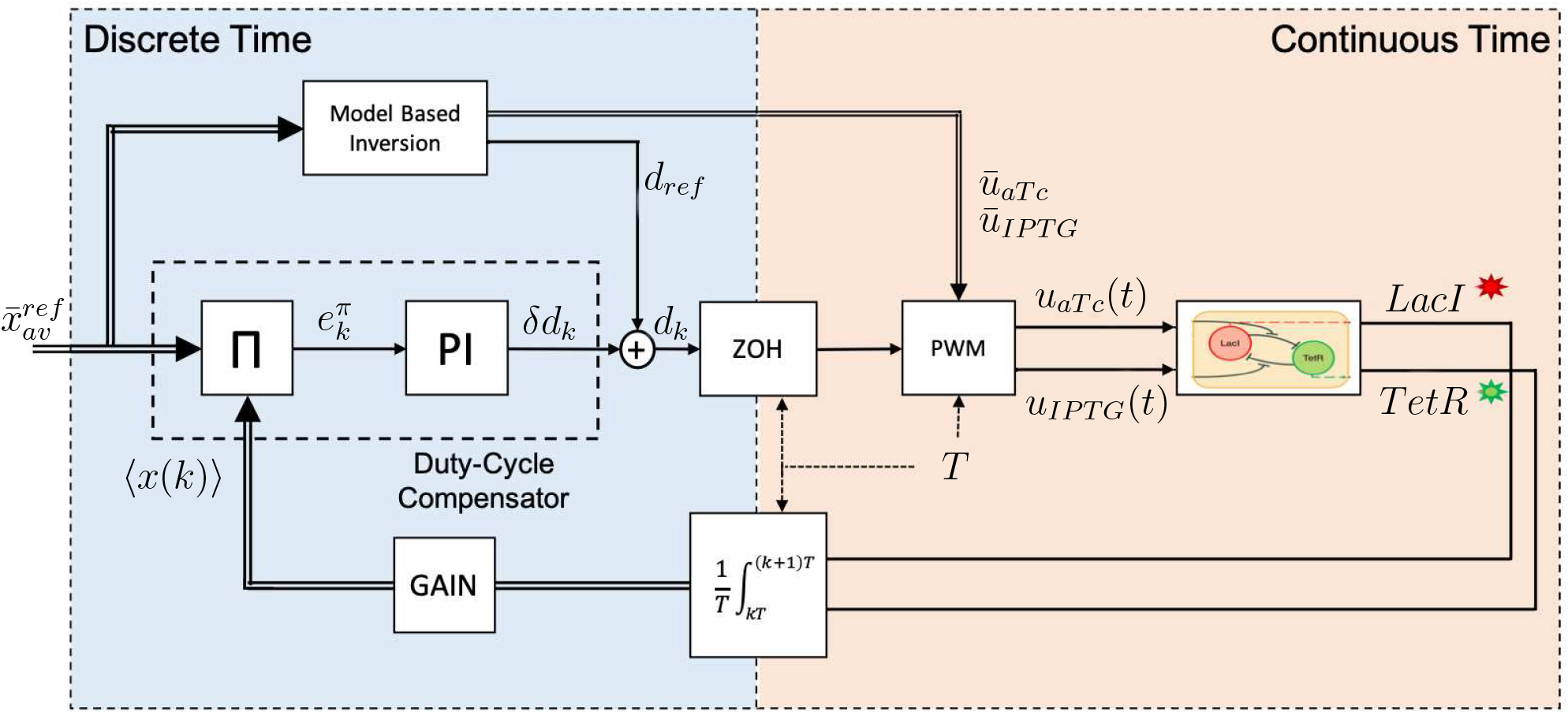
PI-PWM block diagram: Given the setpoint for the average model 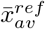, two actions regulate the parameters of the PWM inputs that feed the system. The feedforward action is composed by the Model Based Inversion that evaluates the amplitudes *ū*_*aTc*_ and *ū*_*IPTG*_ and the nominal value of the duty-cycle *d*_*ref*_. The nonlinear projector Π and a proportional-integral controller compose the feedback loop. At each time period *t*_*k*_ = *k T*_*PWM*_, the nonlinear projector Π evaluates the projection error 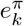 that is minimized by a PI controller that evaluates the correction *δd*_*k*_ to be added to *d*_*ref*_.

These curves of equilibria are exploited in the design of our control strategies, as it is explained later int this text in Sec. 4 and in Sec. 5. For this purpose, we used the average model in Sec. 2 to assemble a database of 60 curves of equilibria, Γ_*i*_(*ū*_*aTc*_, *ū*_*IPTG*_) with *i* = 1, …, 60, parametrized in *d*. The curves were obtained by considering 20 uniformly spaced values of *ū*_*aTc*_ ∈ [0, 100] and 20 values of *ū*_*IPTG*_ ∈ [0, 1]. Specifically, 20 curves were obtained by varying *ū*_*IPTG*_ while keeping constant *ū*_*aTc*_ = 100, 20 curves by varying *ū*_*aTc*_ with *ū*_*IPTG*_ = 1 and 20 by varying simultaneously *ū*_*aTc*_ and *ū*_*IPTG*_ while keeping their ratio constant.

### 4 PI-PWM

The PI-PWM control strategy (*11*, *12*) evaluates the duty-cycle *d*_*k*_ as the sum of *d*_*ref*_ and *δd*_*k*_. The first term, *d*_*ref*_, is the nominal duty-cycle evaluated by using the average model – that neglects the diffusion effects – and depends only on the setpoint value. The second term, *δd*_*k*_, is computed dynamically online by a PI controller whose output is such that the error measure 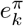, evaluated at the time instant *t*_*k*_ = *k T*_*PWM*_ with a nonlinear projector, decreases.

The feedforward Model Based Inversion works by interrogating a database of 60 curves of equilibria obtained for different values of *ū*_*aTc*_ and *ū*_*IPTG*_ (see Sec. 3). Given the setpoint 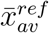, it computes its closest point in the database of equilibria. That point is associated to the values of *ū*_*aTc*_ and *ū*_*IPTG*_ and to a value of the duty-cycle *d*_*ref*_ that are used to initiate the control. This value of the duty-cycle is, then, adapted through the feedback loop, that selects *δd*_*k*_ in real-time.

The core of the PI-PWM algorithm is the nonlinear projector Π that, given the measurements of the mean state value over a time period *T* = *T*_*PWM*_, computed as

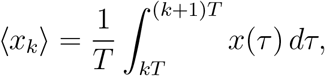

evaluates how far its projection on the closest curve of equilibria 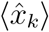 is from the projection 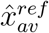 of the setpoint 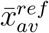. More specifically, as depicted in Figure 10, considering the target equilibrium point of the average model, say 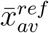, and its current state at time instant *t_k_* = *k T*_*PWM*_, ⟨*x_k_*⟩, the projector Π evaluates their projections on the equilibrium curve 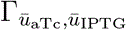, respectively 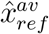 and 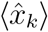. The error 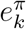 is computed as the arclength between 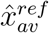 and 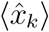 on the curve.

**Figure 10:**
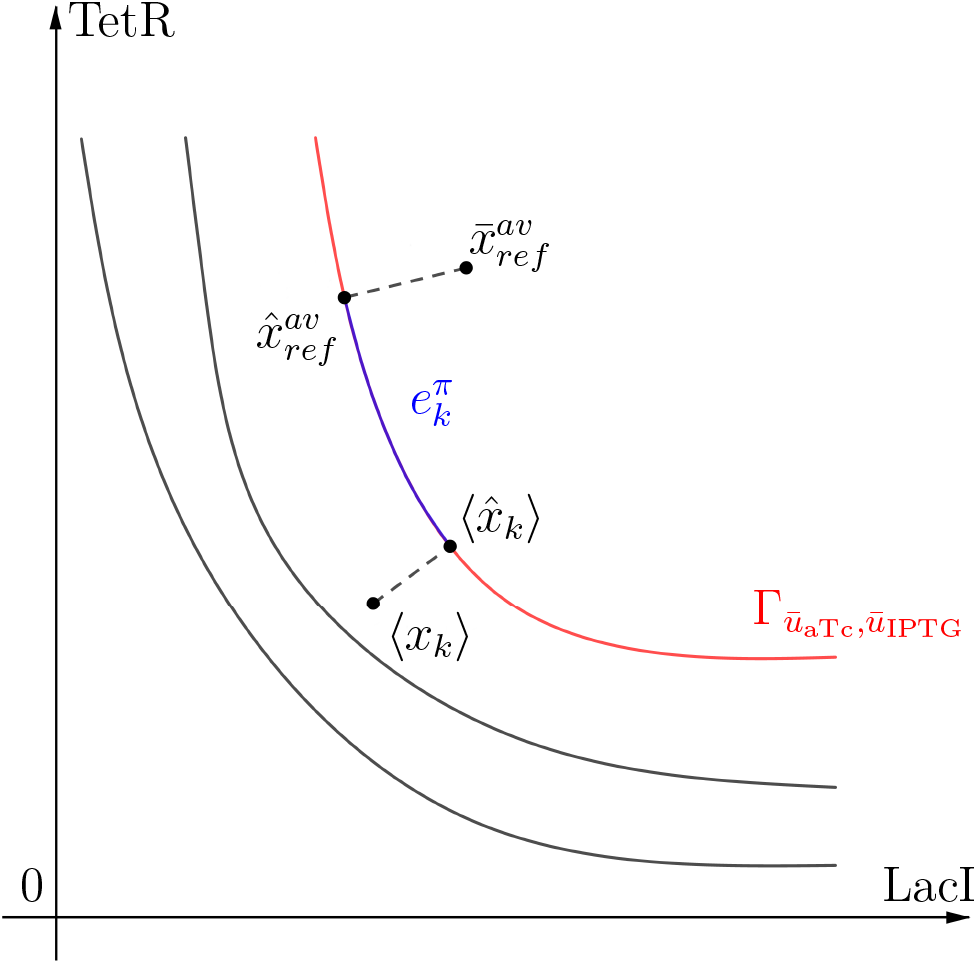
<u>Working principle of the nonlinear projector block</u>. Given the setpoint 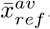, the red curve represents the closest one 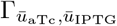; black curves are other equilibrium curves that are further from it. The setpoint 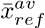 and the mean value of the state in the *k*-th period ⟨*x_k_*⟩ are respectively projected onto the curve on the points 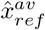 and 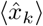. The length of the curve between 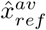 and 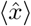, highlighted in blue, is the projection error 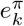 at the time instant *k*.

Note that the projected error 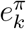 being equal to 0 does not necessarily correspond to zero regulation error of the mean state value ⟨*x_k_*⟩, that is 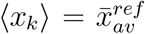. Indeed, at steady-state the line connecting these two points will be orthogonal to the curve 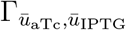 but its length may not be zero. This residual error at steady-state can be made smaller by increasing the number of curves 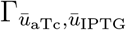 in the database.

The error 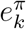, then, is used by a discrete-time proportional-integral PI controller that evaluates the correction *δd*_*k*_ to the duty-cycle at each period as:

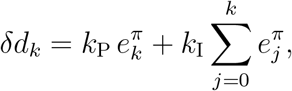

so that the duty-cycle *d*_*k*_ is then set to *d*_*k*_ = *d*_*ref*_ + *δd*_*k*_, starting from *d*_0_ = *d*_*ref*_. This value *d*_*k*_ is then kept constant until the next time instant *t_k_*_+1_ = (*k* + 1)*T*_*PWM*_.

Tuning of the PI gains was carried heuristically via numerical simulations in MATLAB. Specifically, the closed loop system was simulated for 50 periods for 40,000 pairs of gain values *k*_*P*_ and *k*_*I*_ selected uniformly in the ranges *k*_P_ ∈ [10^−4^, 1] and *k*_I_ ∈ [10^−5^, 0.1]; both intervals were divided in 200 uniformly distributed samples. Fig. 11 shows the value of the settling time of the duty-cycle *d*_*k*_ and the norm of steady-state projected error 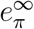 for each pair of gain values. The values of *k*_*P*_ = 0.0101 and *k*_*I*_ = 0.0401 were selected as those giving the best compromise between speed of the transient and residual steady-state error.

**Figure 11:**
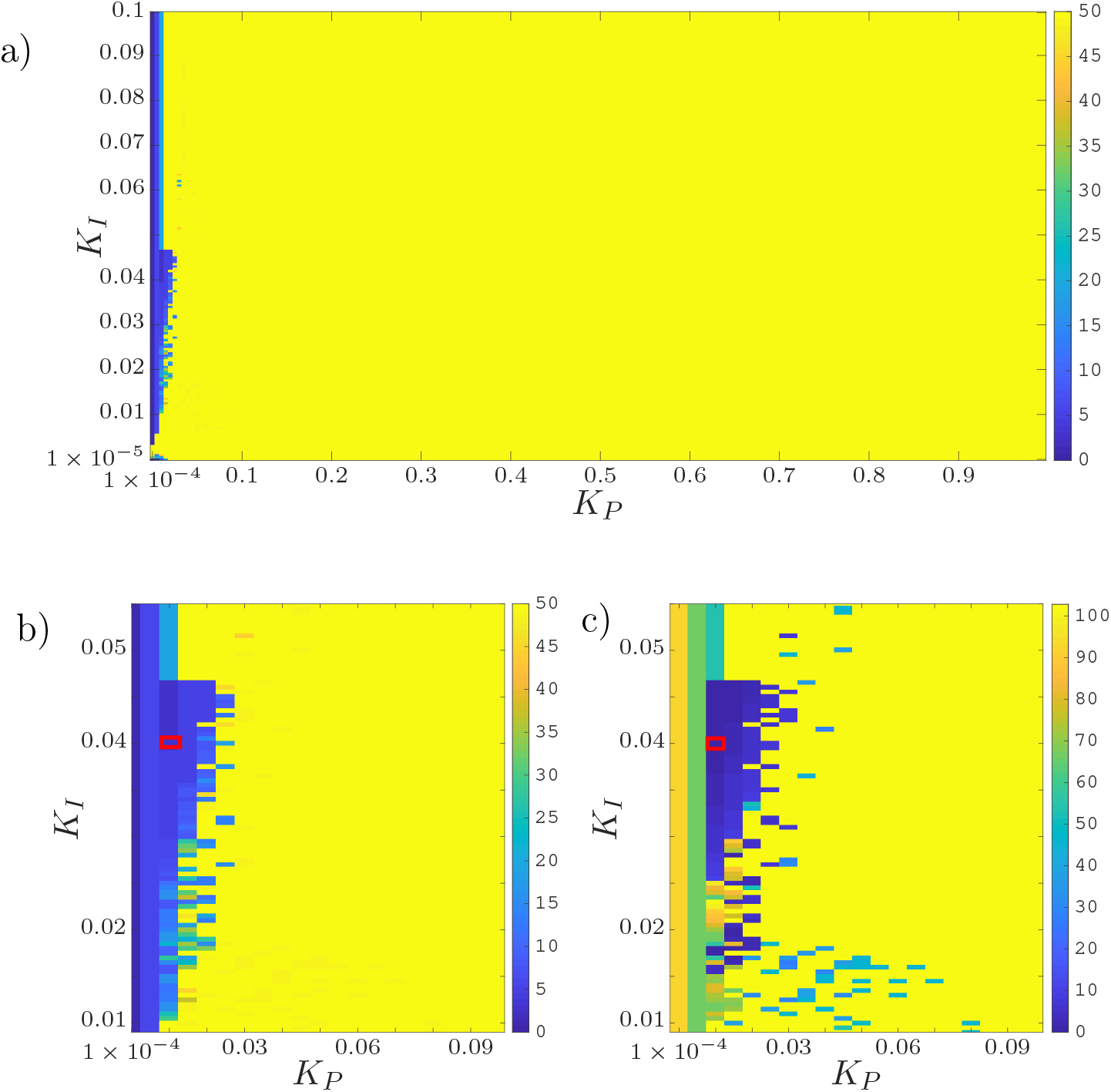
<u>Tuning of the PI controller</u>. Panel a) shows the settling time, as a number of periods, of the duty-cycle at the 10% of its final value, for all pairs (*k*_P_, *k*_I_) ∈ [10^−4^, 1] × [10^−5^, 0.1]. Note that the performance have been evaluated over simulation time of 50 periods and yellow colored squares denote a settling time ≥ 50 periods. Panel b) shows a zoom in the most significant part of panel a) (lower values are better); panel c) shows the norm of the steady-state projected error 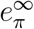 for the same range of values of control gains (lower values are better). The red box indicates the values of PI gains we selected and used in all in-silico control experiments.

**Figure 12:**
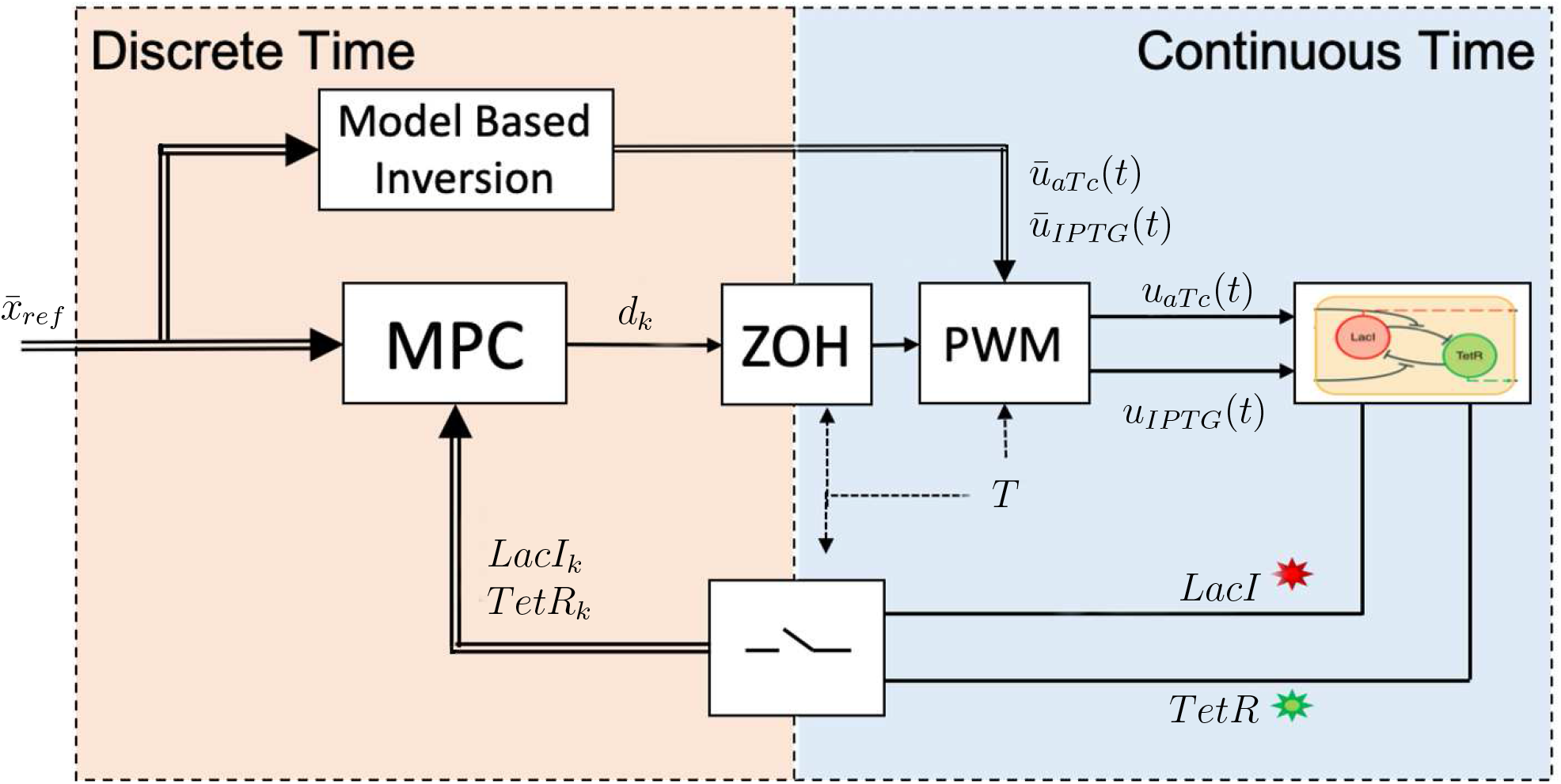
Model predictive control block diagram

### 5 Model Predictive Control

MPC strategies have been widely applied in the field of synthetic biology (*29*), demonstrating their effectiveness also for in-vivo experiments (*22*). A Model Predictive Controller repeatedly solves an online optimization problem (on a finite prediction horizon interval) to directly evaluate the duty-cycle *d*_*k*_ as the quantity that minimizes a given cost function; in our case the cost function was selected as:

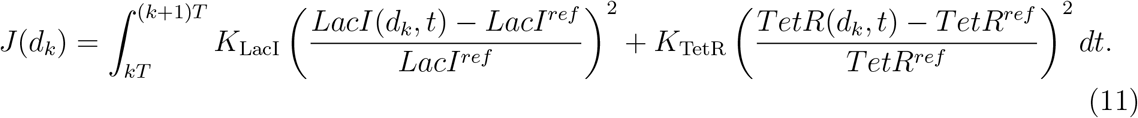

The optimization problem is formulated taking into account the 6^*th*^ order model of the system (see Sec. 1 of the SI, equations (1)–(6)) to predict its state evolution over an horizon that is multiple of *T*_*PWM*_, that is *T*_*p*_ = *n*_*ph*_ · *T*_*PWM*_ with 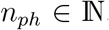. Other constraints, as the upper and the lower admissible bounds of the duty-cycle, were considered in the formulation of the problem. Genetic algorithms (*30*) were used to numerically find the (sub)optimal solution at each step.

The control parameters *K*_LacI_ and *K*_TetR_ in (11) were selected heuristically to *K*_LacI_ = 1 and *K*_TetR_ = 4, after an extensive numerical search in MATLAB. Specifically, the control evolution was simulated for 18 periods fixing *K*_LacI_ = 1 and varying *K*_TetR_ over 37 values chosen uniformly in the interval [0.01, 100], so as to vary the ratio between the two gains. The best values given above were selected for the in-silico experiments reported in the main text.

### 6 Comparison Metrics

To conduct a quantitative comparative analysis between different control approaches, we evaluated three different control error metrics as done in (*22*). The Integral Square Error (ISE) is defined as

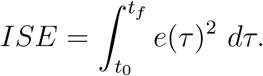

By integrating the square error over time, ISE tends to penalize large errors much more than smaller ones. Thus, this index can be used to compare performance during transients, in which the presence of the overshoot or a long settling time could give rise to significant errors. Small errors, even if persistent, do not affect this metric in a very significant way. The Integral Absolute Error (IAE) is defined as

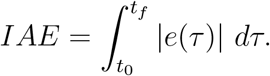

IAE integrates the absolute error without adding any weight. This index allows a trade-off between transient dynamics and steady-state errors.

The Integral Time-weighted Absolute Error (ITAE) is defined as

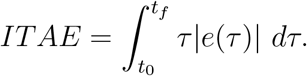

By integrating the absolute error multiplied by the time, ITAE tends to penalize much more small errors that occur after a long time than significant errors at the beginning of the experiment. For this reason, it is the best index to summarize the performance of a controller in a regulation task.

In the above equations, *t*_0_ and *t*_*f*_ are, respectively, the time instants at the beginning and at the end of the in-silico experiments.

### 7 Quantitative analysis of the in-silico experiments

Performance indices of the deterministic in-silico experiments are reported in Table 2. The comparison metrics, presented in the previous Sec. 6, have been evaluated by using the relative error signal *e*(*t*) defined as

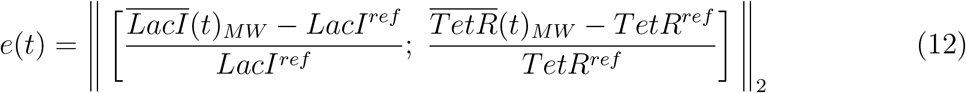

where

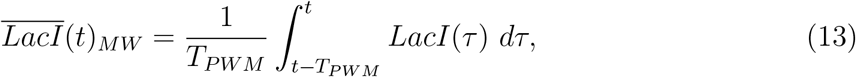

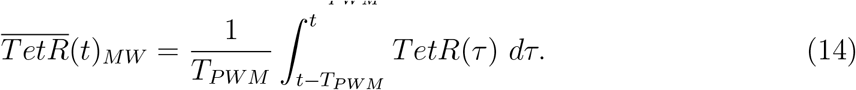

**Table 2:** Control Error Metrics in the *deterministic* case. The two strategies are compared in terms of *Integral Square Error* (ISE), *Integral Absolute Error* (IAE), and *Integral Time-weighted Absolute Error* (ITAE). The error signal is computed according to (12).

| Strategy | ISE | IAE | ITAE |
| --- | --- | --- | --- |
| PI-PWM | 876.71 | 1304.32 | 2.068673 E06 |
| MPC | 47.58 | 382.04 | 0.811109 E06 |

They confirm the qualitative analysis reported in the main text. Generally, MPC leads to better performance than PI-PWM in the deterministic case, i.e. where stochastic perturbations are not taken into account.

In the stochastic experiments, we find very similar results as in the deterministic case. MPC still globally guarantees better performances, even if its indices worsened more than those computed for the PI-PWM (Table 3).

**Table 3:** Control Error Metrics in the *stochastic* case. The open loop forcing and the two control strategies are compared in terms of *Integral Square Error* (ISE), *Integral Absolute Error* (IAE), and *Integral Time-weighted Absolute Error* (ITAE) of the error signal, computed according to (12) over the last 12 periods. The values reported in brackets are computed over the entire simulation (18 periods).

| Strategy | ISE | IAE | ITAE |
| --- | --- | --- | --- |
| Open Loop | (13456.03) | (7413.06) | (16.822305 E06) |
| PI-PWM | 469.90 (830.52) | 998.20 (1472.87) | 2.903351 E06 (3.218139 E06) |
| MPC | 150.14 (178.50) | 616.05 (724.72) | 1.904908 E06 (1.975185 E06) |

#### Parametric variations

To test robustness of the strategies, control error metrics were computed in stochastic simulations with parametric variations and are reported in Table 4. The values reported in the table are evaluated by averaging three simulations each obtained using different sets of parameters.

**Table 4:** Control Error Metrics in the *stochastic* case with *parametric variations*. The parameters of each cell in the population were independently drawn from Gaussian distributions centered on their nominal values given in Tab. 1 (in Section 1) with standard deviation of 5% and 10%, respectively. The open loop forcing and the two control strategies are compared in terms of *Integral Square Error* (ISE), *Integral Absolute Error* (IAE), and *Integral Time-weighted Absolute Error* (ITAE) of the error signal, which has been computed according to (12) over the last 12 periods of the simulations. Values reported in brackets are related to the entire simulations (18 periods).

| Strategy | Var. | ISE | IAE | ITAE |
| --- | --- | --- | --- | --- |
| Open Loop | 5% | (15613.06) | (7985.07) | (18.091973 E06) |
| PI-PWM | 5% | 687.30 (1605.77) | 1224.34 (2056.69) | 3.605903 E06 (4.162760 E06) |
|  | 10% | 4369.72 (6061.33) | 3094.09 (4317.68) | 9.014108 E06 (9.968271 E06) |
| MPC | 5% | 1116.21 (1459.63) | 1532.02 (2109.08) | 4.193415 E06 (4.716253 E06) |
|  | 10% | 2185.18 (2751.82) | 2164.75 (2870.43) | 6.153388 E06 (6.791182 E06) |

In the case of parameters extracted with a standard deviation of 5%, MPC guarantees better transient performance – as confirmed by lower ISE value over the whole simulation – while PI-PWM achieves better steady-state regulation, i.e. lower ITAE values. While in the case of a standard deviation of 10%, PI-PWM metrics worsen more than for the MPC, which globally guarantees better performance.

#### Agent-based simulations in BSim

Fig. 13 shows the evolution of the cells’ output when simulated in open-loop (i.e. without feedback control action) via agent-based simulations. When comparing this figure with the closed-loop simulation shown in the main text (Fig. 6) we see that LacI and TetR average values are consistently far away from the reference setpoints, as opposed to their evolution in the closed-loop simulations.

**Figure 13:**
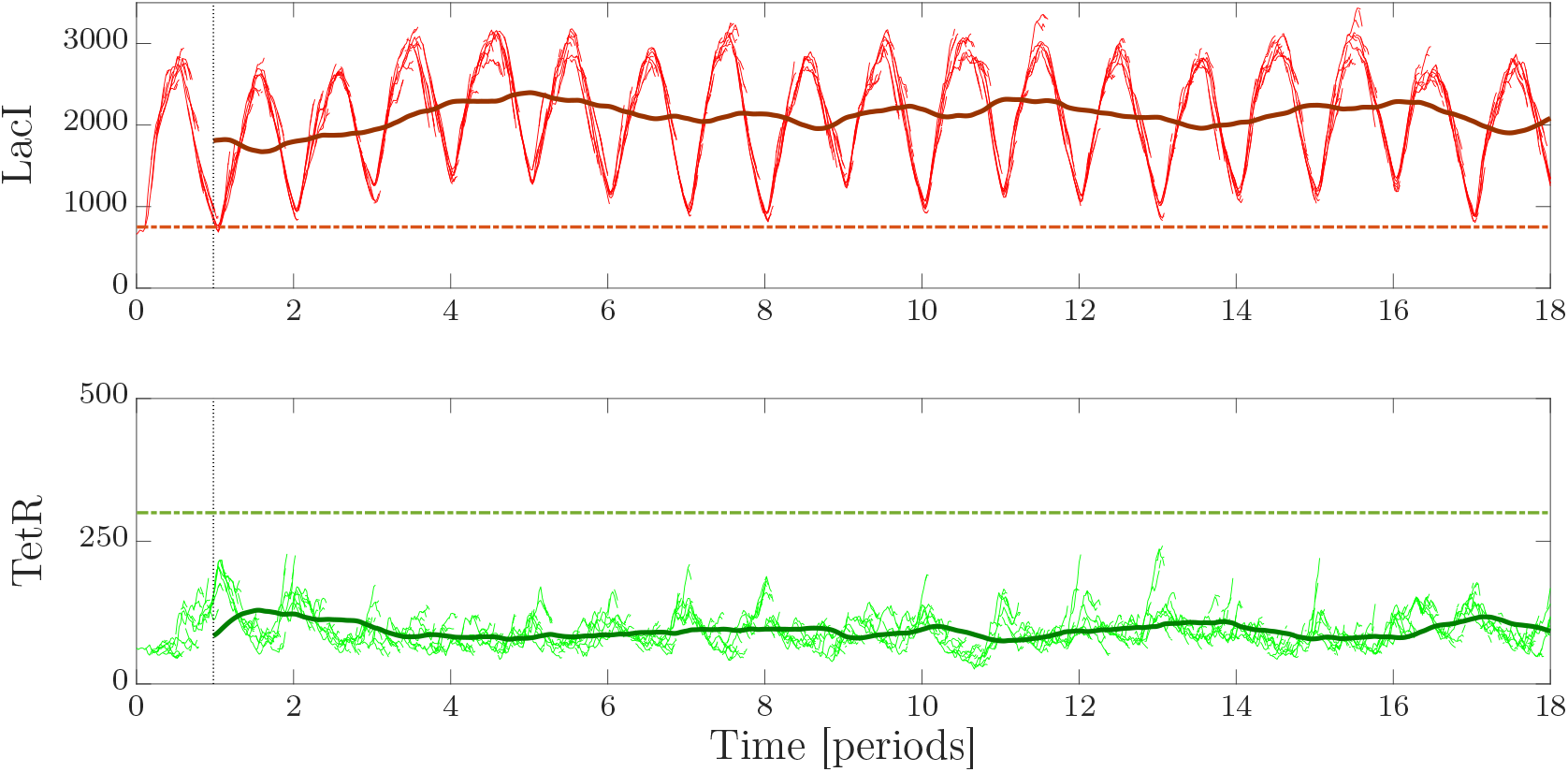
<u>Agent-based simulation in BSim of the cells evolution in open-loop</u>. The pulsatile inputs’ amplitudes were set to *ū*_*aTc*_ = 35, *ū*_*IPTG*_ = 0.35, while the duty-cycle was kept constant (without any adaptation) to *d*_*ref*_ = 0.4. The period was selected as usual to be *T*_*PWM*_ = 240 min. Total simulation time is 72 hours. We considered E. coli cells growing in a single chamber of a “mother machine”-like microfluidic device (*27*): the simulations start with a single cell located at the bottom of the chamber; as the cell grows and duplicates, it pushes outside of the chamber new cells that exceed the maximum capacity of the chamber (around 10 cells). The top panel shows the evolution over time of *LacI*; the dashed line representing the setpoint *LacI*_*ref*_ = 750, while lighter lines the evolution of the state for each cell in the simulation, and the darker solid line the mean trajectory computed over the population, evaluated through a moving window of period *T*_*PWM*_. The middle panel shows the evolution over time of *TetR*; the dashed line representing the setpoint *TetR*_*ref*_ = 300, lighter lines are the evolution of the state for each cell in the simulation, while the dark solid line represents the evolution of the mean trajectory across the population in the period, evaluated using a moving window of period *T*_*PWM*_.

Variations of the 10% on all the parameters of the system do not influence the overall performance of the two closed-loop control strategies, with the MPC still considerably better than the PI-PWM (see Table 5).

**Table 5:** Control Error Metrics in the agent-based simulations. The open-loop forcing and the proposed control strategies are compared in terms of *Integral Square Error* (ISE), *Integral Absolute Error* (IAE), and *Integral Time-weighted Absolute Error* (ITAE) of the error signal that is computed according to (12). The values are computed neglecting the transient of about 5 periods required for the circuit outputs to settle down. The values computed over the entire simulation are reported in brackets.

| Strategy | Var. | ISE | IAE | ITAE |
| --- | --- | --- | --- | --- |
| Open Loop | no | 11161.13 (15543.92) | 5661.29 (7923.19) | 16.271114 E06 (18.281741 E06) |
| PI-PWM | no | 1483.72 (3671.39) | 1510.72 (2939.98) | 3.452511 E06 (4.458611 E06) |
|  | 10% | 1781.40 (3508.97) | 1826.95 (2975.92) | 4.477333 E06 (5.217549 E06) |
| MPC | no | 173.12 (277.18) | 562.83 (836.80) | 1.677061 E06 (1.947358 E06) |
|  | 10% | 283.49 (481.36) | 732.75 (1201.13) | 2.097497 E06 (2.451896 E06) |

## Footnotes

1 https://it.mathworks.com/matlabcentral/fileexchange/34707-gillespie-stochastic-simulation-algorithm

2 https://it.mathworks.com/help/gads/ga.html

